# Cross-species cell-type assignment of single-cell RNA-seq by a heterogeneous graph neural network

**DOI:** 10.1101/2021.09.25.461790

**Authors:** Xingyan Liu, Qunlun Shen, Shihua Zhang

## Abstract

Cross-species comparative analyses of single-cell RNA sequencing (scRNA-seq) data allow us to explore, at single-cell resolution, the origins of cellular diversity and the evolutionary mechanisms that shape cellular form and function. Here, we aimed to utilize a heterogeneous graph neural network to learn aligned and interpretable cell and gene embeddings for cross-species **c**ell type **a**ssignment and gene **m**odule **e**xtraction (CAME) from scRNA-seq data. A systematic evaluation study on 649 pairs of cross-species datasets showed that CAME outperformed six benchmarking methods in terms of cell-type assignment and model robustness to insufficiency and inconsistency of sequencing depths. Comparative analyses of the major types of human and mouse brains by CAME revealed shared cell type-specific functions in homologous gene modules. Alignment of the trajectories of human and macaque spermatogenesis by CAME revealed conservative gene expression dynamics during spermatogenesis between humans and macaques. Owing to the utilization of non-one-to-one homologous gene mappings, CAME made a significant improvement on cell-type characterization cross zebrafish and other species. Overall, CAME can not only make an effective cross-species assignment of cell types on scRNA-seq data but also reveal evolutionary conservative and divergent features between species.

## Introduction

Single-cell RNA sequencing (scRNA-seq) has rapidly emerged as a powerful tool to characterize a large number of single-cell transcriptomes in different tissues, organs, and species [2]. It not only deepens our knowledge of cells but also provides novel insights into evolutionary and developmental biology [3]. Cross-species integration and comparison of scRNA-seq datasets allow us to explore, at single-cell resolution, the origins of cellular diversity and the evolutionary mechanisms that shape cellular form and function [3-11].

Cell-type assignment (or cell typing) and data integration are both vital steps involved in these analyses. For the cell-type assignment, a traditional approach includes three steps: clustering single-cells, performing differentially expression analysis to find cluster-specific genes, and matching these genes with known markers. However, this strategy fails when different cell types are clustered into one group, and when analyzing many non-model species that lack prior knowledge of cell-type biomarkers. Several tools have been developed for this task recently. Some existing approaches like CellAssign [12] and scCATCH [13] require prior knowledge of cell type-specific markers. Some like SingleCellNet [14] and SciBet [15] were designed based on a reference dataset and can achieve the cell-type assignment without providing marker information. Besides, several methods designed for data integration can also achieve cell-type assignment by transferring labels from the reference dataset. Seurat-v3 [16] combines canonical correlation analysis and mutual nearest neighbors to perform data integration and label transfer based on ‘anchors’. Cell BLAST [17] and ItClust [18] make use of deep neural networks for both cell-type querying and cell embedding. LIGER [19] and CSMF [20] extract the common and private features of two datasets respectively by joint non-negative matrix factorization to achieve cell alignment across datasets and omics.

Despite all the progress, a tool for effective and robust cross-species integration and comparison is still immature and in demand. There are several computational challenges to be overcome. First, it is hard to determine cell identities for non-model species that lack prior knowledge of cell-type biomarkers, and most of the methods may fail when generalizing to cross-species label transfer. Second, many biological and technical factors, such as transcriptome variation between species, different experimental protocols, and inconsistent sequencing depths, can make cross-species data integration and comparison even more difficult. Third, homologous cell-type alignment requires quantifying the similarities of gene expression profiles, which usually vary across distinct normalizations and gene selections [3]. Fourth, cross-species cellular alignment is usually based on homologous genes and current approaches are mostly restricted to one-to-one homologies shared by both organisms [3, 5-11], where non-one-to-one homologous genes characterizing cell-type conservative features could be lost. Lastly, evolutionary divergences are thought to be caused by transcriptional changes of groups of genes that evolve in a modular fashion and are controlled by transcription factors [21]. Extraction and comparison of gene modules between species will provide deep insights into evolutionary conservation and divergences [11, 22, 23].

To this end, we developed a heterogeneous graph neural network model to achieve the aligned and interpretable cell and gene embeddings for cross-species **c**ell-type **a**ssignment and gene **m**odule **e**xtraction (CAME). A systematic evaluation study on 649 pairs of cross-species datasets showed that CAME outperformed six benchmarking methods in terms of cell-type prediction, and model robustness to insufficiency and inconsistency of sequencing depths. Comparative analyses of the major types of human and mouse brains by CAME revealed shared cell type-specific functions in homologous gene modules. An alignment of the trajectories of human and macaque spermatogenesis by CAME revealed the conservative gene expression dynamics during spermatogenesis between humans and macaques. Owing to the utilization of non-one-to-one homologous gene mappings, CAME made a significant improvement on cell-type characterization across long-distant species. Overall, CAME can not only make an effective cross-species assignment of cell types on scRNA-seq data but also reveal evolutionary conservative and divergent features between species.

## Results

### Overview of CAME

CAME takes two scRNA-seq datasets from different species, along with their homologous gene mappings as input. One dataset with cell-type labels is taken as the reference and the other whose cell types need to be assigned is the query (**Figure 1A**). CAME encodes these two expression matrices and the mappings of homologous genes as a heterogeneous graph, where each node acts as either a cell or a gene, while a cell-gene edge indicates a non-zero expression of the gene in that cell, and an edge between a pair of genes indicates the homology between each other. Note that one-to-many and many-to-many homologies are allowed as well. Besides, CAME adopts single-cell networks pre-computed from reference and query datasets using the k-nearest-neighbor (KNN) method, respectively, where a cell-cell edge indicates this pair of cells have similar transcriptomes with each other (**Methods**).

**Figure 1.**
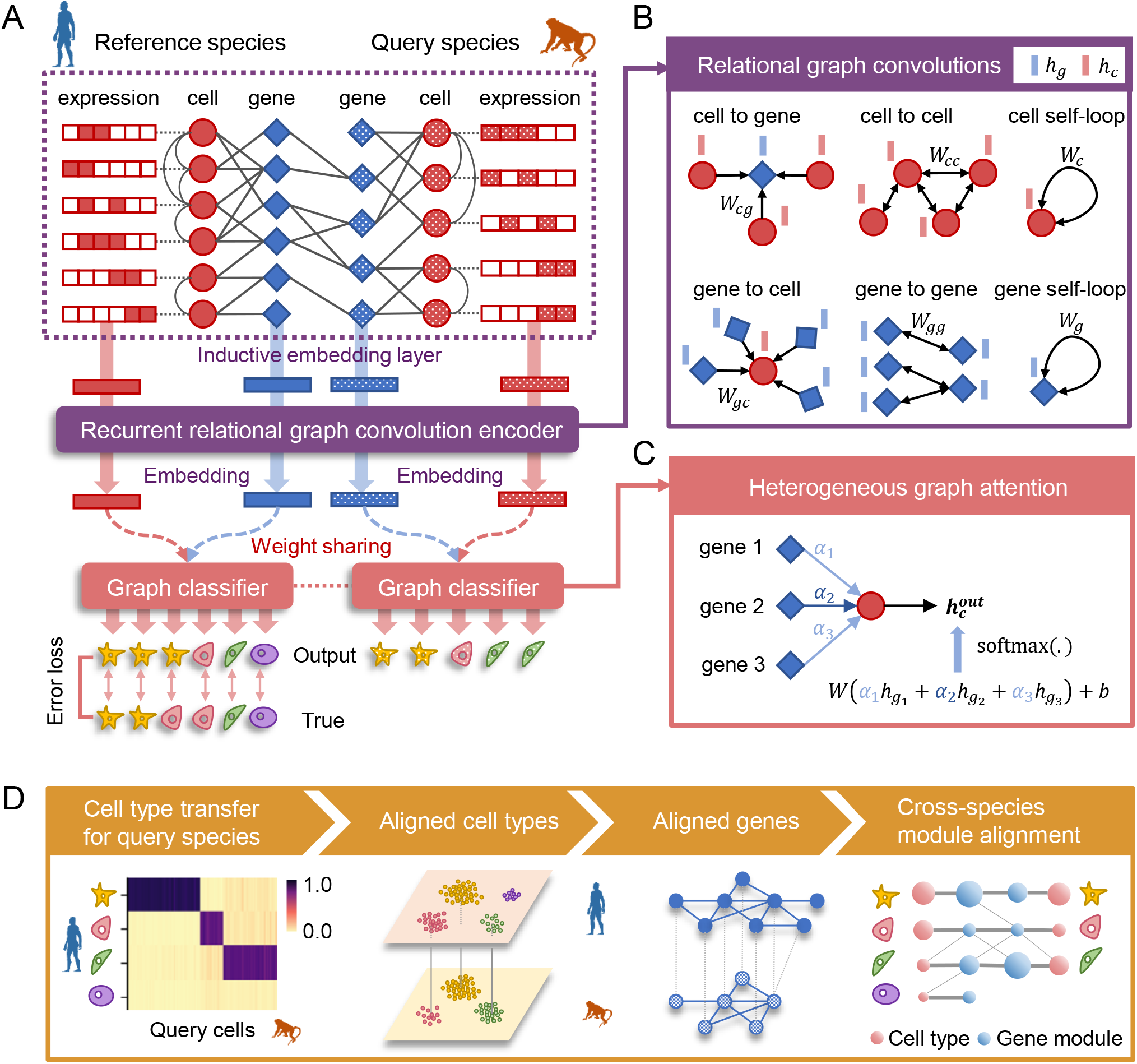
Overview of CAME. (**A**) The architecture of the heterogeneous graph neural network in CAME. The scRNA-seq data of both reference and query species and their homology genes are encoded as a heterogeneous cell-gene graph. The cell-gene edge indicates that the cell has non-zero expression of the gene. The gene homologous mappings are represented by a gene-gene bipartite graph with each edge indicating a gene homology. Note that the homologous gene mappings can be many-to-many homologies. To preserve the intrinsic data structure, the within-species cell-cell edges are adopted where an edge between a pair of cells indicates that one is the k nearest neighbor of the other (k=5 by default). The heterogeneous graph and the gene expression profiles are input to CAME, passing through the inductive embedding layer, the recurrent relational graph neural network, and the graph classifier with attention mechanisms. The model is trained by minimizing the cross-entropy loss calculated between the model prediction and the given labels of the reference cells in an end-to-end manner. (**B**) Graph spatial convolutions for six different types of edges including “a cell expresses a gene”, “a gene is expressed by a cell”, “cell-cell similarity”, “gene-gene homology”, “cell self-loop” and “gene self-loop” with the edge type-specific convolution weights. (**C**) Heterogeneous graph attention classifier on the last layer, where each cell pays different attention to its neighbor genes. The output cell-type probabilities are calculated by the weighted sum of the neighbor-gene embeddings, followed by the softmax normalization. The attention weights are calculated from the concatenated cell and gene embeddings with a linear transformation, followed by activation and the softmax normalization among the neighbor-genes of the cell. (**D**) The output of CAME includes the probabilistic cell-type assignment of the query species, as well as low-dimensional embeddings of the cells and genes from both species. The gene embeddings are used for joint module extraction that allows inter-species comparison of conservative or divergent characteristics.

CAME adopts a heterogeneous graph neural network to embed each node into a low-dimensional space (**Methods, Figure 1B**). For the initial cell embeddings, CAME takes the expression profiles followed by linear transformation with a non-linear activation function. While for the initial gene-embeddings, CAME aggregates the expression profiles (called “message”) from its neighbor cells which expressed it, and then treats them with linear transformation and non-linear activation, as done for cells (**Methods**). Then the initial embeddings are input to two parameter-sharing graph convolution layers with heterogeneous edges and nodes. As a result, cells with more co-expressed genes are more likely to exchange the embedding message with each other, thus be encoded with similar embeddings; the same principle applies to genes. CAME further employs a heterogeneous graph attention mechanism [25] to classify cells with embeddings of their neighbor genes as input, where each cell pays a distinct level of attention to each certain neighbor gene (**Methods, Figure 1B**). High attention paid by a cell to a gene implies that the gene is of relatively much importance for the cell to be characterized.

We note that a reference cell could be assigned with multiple labels in different hierarchies, and a cell type in query species might correspond to multiple ones in the reference. Thus, multi-label classification can be helpful to depict the state of a cell. CAME calculates the cross-entropy between the predicted cell-type probabilities and the true labels for the reference data to obtain both the multi-class and the multi-label loss, and sums them up as the training loss. Finally, CAME minimizes it by the backpropagation algorithm (**Methods**). The training process of CAME is semi-supervised in an end-to-end manner. We found that the training process was quite stable, and the model tended to be well trained before 200-300 epochs (**Supplementary Figure S1A**). Besides, CAME introduces the adjusted mutual information (AMI) between the predicted labels and pre-clustered ones of query cells to automatically determine the model checkpoint for downstream analysis (**Methods** and **Supplementary Figure S1A**). Ablation experiments demonstrated that six key factors adopted by CAME play roles in improving the prediction performance (**Supplementary Figure S1B**).

CAME outputs the quantitative cell-type assignment for each query cell, that is, the probabilities of cell types that exist in the reference species, which enables the identification of the unresolved cell states in the query data. For most cells with homologous cell types in the reference, CAME assigns them with a maximal probability approximating 1. While for those unobserved cell types or states, CAME would assign them to their analogs with relatively low confidences **(Supplementary Figure S2)**. Besides, CAME gives the aligned cell and gene embeddings across species, which facilitates low-dimensional visualization and joint gene module extraction (**Methods, Figure 1D**).

### CAME showed superior accuracy and robustness for cell-type assignment compared to benchmarking methods

We collected 54 scRNA-seq datasets from five tissues across seven different species including human, macaque, mouse, chick, turtle, lizard, and zebrafish (**Methods, Supplementary Figure S3A** and **Supplementary Table S1**) and found that more than a half of the homologous genes between zebrafish and other species are not one-to-one matched (**Supplementary Figure S3B**). Besides, the proportion of non-one-to-one homologies between highly informative gene (HIG) sets with one associated with zebrafish [26] was significantly higher than that of other cross-species dataset pairs (60%-75% versus 15%-40%, **Supplementary Figure S3C**). And ablation study shows that, when excluding non-one-to-one homologies, the cell-typing accuracy of CAME suffered a significant drop (ranging from 1.5% to 8.7% for different species-pairs, 6.26% on average, with p-value = 7.8e-23) on the zebrafish-associated dataset pairs (**Supplementary Figure S3D and Figure S4**). Therefore, we divided these pairs into two scenarios: zebrafish-excluded (139 pairs) and zebrafish-associated (510 pairs) (**Methods**).

We compared the cell-typing performance of CAME with six benchmarking methods including two marker-based methods SciBet [15] and Scamp [44], two deep-learning methods Cell BLAST [17] and ItClust [18], one expression-based method SingleCellNet [14], and one integration-based method Seurat-v3 [16] in these two scenarios in terms of accuracy, macro-F1 score and weighted F1 score (**Methods**). Results showed that, in both scenarios, CAME distinctly outperformed the others in most cases with statistical significance p-values < 10^−16^ and 10^−54^ using Wilcoxon signed-rank test for both zebrafish-excluded and zebrafish-associated scenarios, respectively (**Figure 2A and B, Supplementary Figures S5 and S6**).

**Figure 2.**
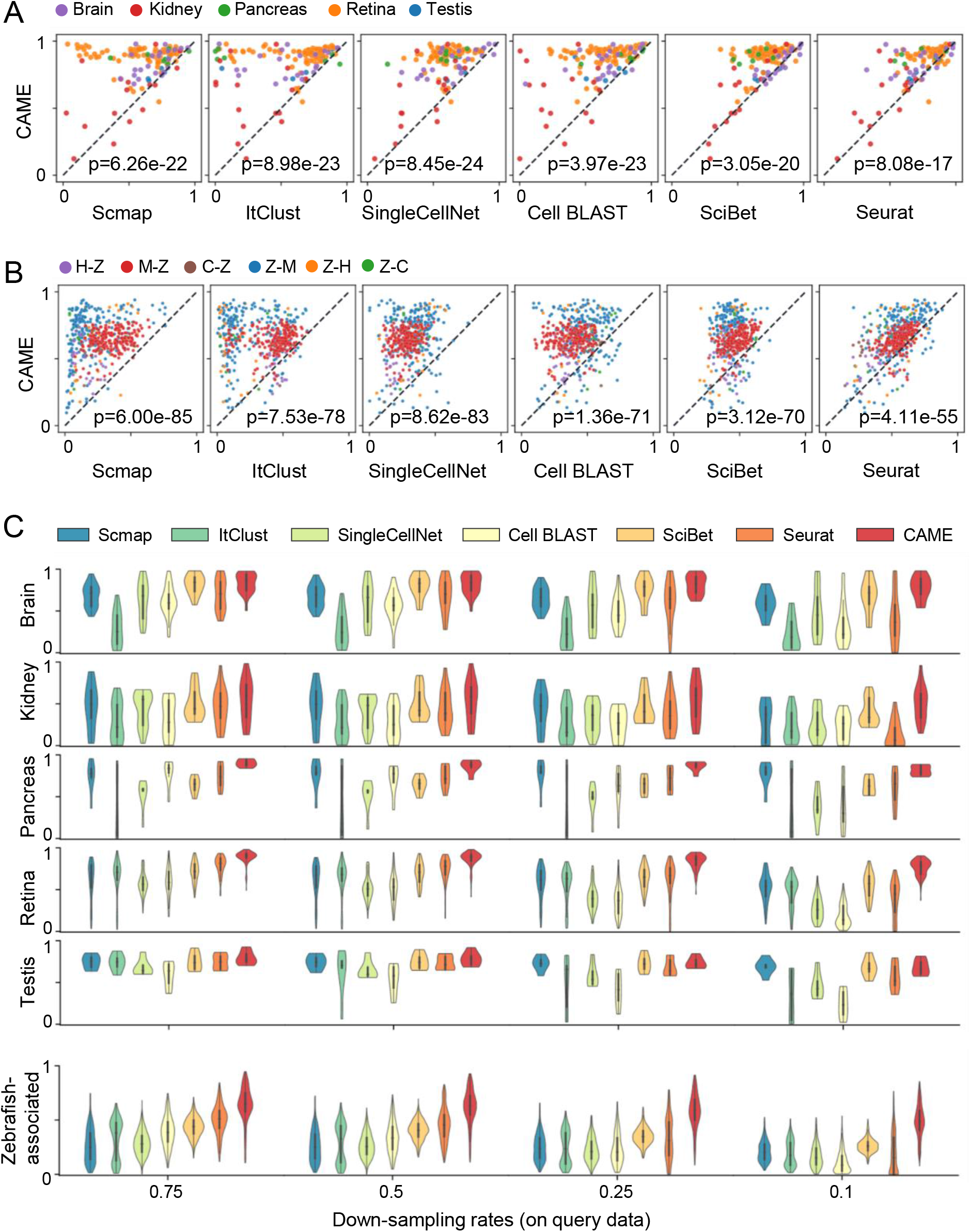
Benchmarking cross-species cell-type assignment performance of CAME. (**A** and **B**) Performance comparison of CAME and the six benchmarking approaches in terms of cell-typing accuracy on 139 pairs of cross-species scRNA-seq datasets (A) and on 510 pairs of cross-species scRNA-seq datasets that associated with zebrafish, where each point represents a pair of cross-species datasets and is colored by tissue. The notation “X-Y” indicates that X is the reference and Y is the query. H: Human, M: Mouse, C: Chick, Z: Zebrafish. (C) Performance comparison of the classification accuracies of CAME and the six benchmarking methods on different down-sampling rates (0.75, 0.5, 0.25, 0.1) for read counts.

To evaluate the robustness of CAME in the cases when the reference and query datasets have inconsistent and insufficient sequencing depths, we performed down-sampling experiments (at various sampling rates 75%, 50%, 25%, 10%) for read counts on the reference, query, and both reference and query datasets. Again, CAME achieved superior performance compared to all six benchmarking methods (**Figure 2C, Supplementary Figures S7 and S8**). By contrast, when the down-sampling rates are extremely unbalanced, some benchmarking methods may fail. For example, at a down-sampling rate of 0.1 for query datasets, Seurat detected too few anchors to abort integration for label transfer and Scmap failed to find enough genes since the median expression in the selected features is 0 in each cell cluster. All these results demonstrate that CAME is robust to the insufficient and inconsistent sequencing depths between reference and query pairs.

### CAME could robustly align homologous cell types across species and multiple references

In addition to the accurate cross-species cell-type assignment, CAME is also capable of aligning homologous cell types from different species, even when crossing distant species. For example, when aligning cell types between mouse [29] and turtle [10], CAME successfully distinguished and aligned each major type, like inhibitory and excitatory neurons, while the alignments by FastMNN, Harmony, and Seurat were incapable. CAME also separated the neural progenitor cells from excitatory neurons, while LIGER merged these two groups. The visualization plots using Uniform Manifold Approximation and Projection (UMAP) [31] of cell embeddings of Cell BLAST tend to lose some relations between cell types, e.g., the inhibitory and excitatory neurons are not linearly separable on the 2D plot (**Figure 3A**, and **Supplementary Figure S9**).

**Figure 3.**
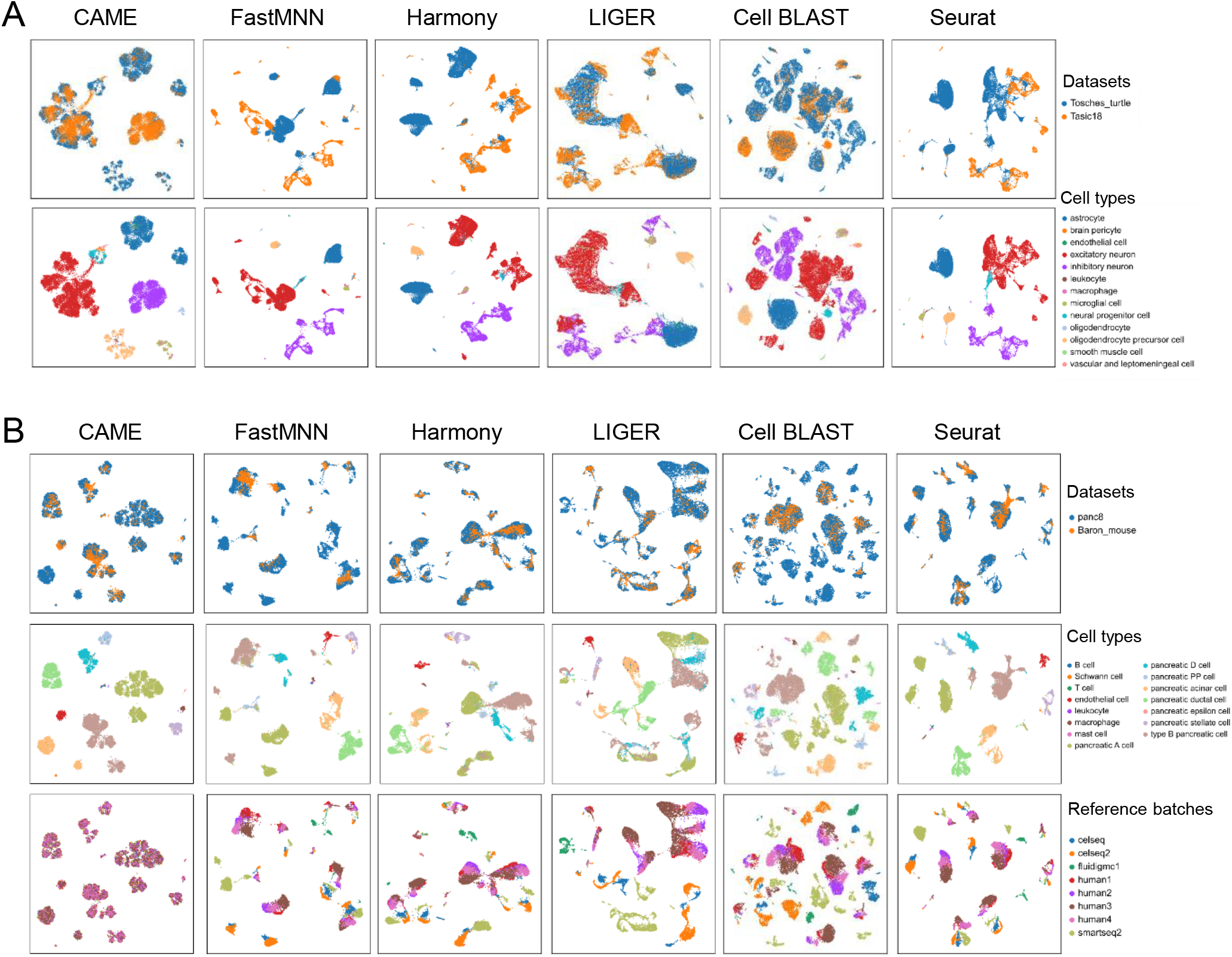
Alignment comparison of cell embeddings across datasets by CAME and five benchmarking methods. (**A**) The UMAP plots of the cell embeddings by CAME and five benchmarking integration methods on the scRNA-seq data from turtle (reference) and mouse (query) brains. Cells are colored by their dataset identities (the first row) or cell type (the second row). (**B**) Similar settings to (A). Here the reference datasets are the human pancreatic scRNA-seq data from eight batches by five different platforms and the query is from mouse pancreas cells. The UMAP plots of the third row showed the reference datasets, colored by batch identities.

When handling multiple references and batch information is unavailable, most integration methods will suffer from batch effects. In this situation, owing to the semi-supervised manner, CAME can ignore the batch effects of reference data. In contrast, other integration tools may suffer from diverse sources of noises if the potential batch effects (such as noises from different individuals) are not considered. For instance, when aligning human and mouse pancreas cell types with human reference composed of eight batches, cells of the same type but from different batches were still separated from each other. Besides, the query cells tended to be “attracted” by reference cells of the same protocol (**Figure 3B**). Even when the batch labels are given, for some of the benchmarking methods (e.g., LIGER [19] and Seurat-v3 [16]), the reference batch effects still existed after data integration (**Supplementary Figure S10**).

### CAME could accurately assign cell types in mouse brains and reveal cell-type-specific gene modules

We applied CAME to assign the major types of single cells from the primary visual cortex and the anterior lateral motor cortex of mice [29], and used human brain cells as the reference dataset [7], containing the cells from the hindbrain that is not included in the mouse dataset. CAME achieved an accuracy of about 98%, so as Seurat and SciBet, superior to other benchmarking methods (94% by ItClust, 93% by Cell BLAST, 92% by SingleCellNet, and only 55% by Scmap). CAME also got a higher macro-F1 score (0.55) than that of Seurat (0.44) and SciBet (0.46), indicating that CAME also accurately classified the small groups. Specifically, those non-neuronal types accounting for a small proportion of mouse cells were accurately assigned, including endothelial cells (accounting for 0.6% of human cells and 0.85% of mouse cells) and its subclass, brain pericytes (0.61% of human cells and 0.14% of mouse cells). The macrophages (0.56% in mice) were classified as microglial cells (2.1% of human cells) that are biologically similar to this type. Both oligodendrocyte precursor cells (OPC) and oligodendrocytes in mice were originally assigned as oligodendrocytes (0.75% of mouse cells) by the authors, but they were distinguished from each other in the reference of the human data (**Figure 4A**). The identities of OPCs were also verified by examining the expression of typical marker genes in each cell type (**Figure 4B**). Besides, we found that the genes with top attentions from each cell type showed high cell-type specificities, though these genes were quite different across species (**Supplementary Figure S11A**).

**Figure 4.**
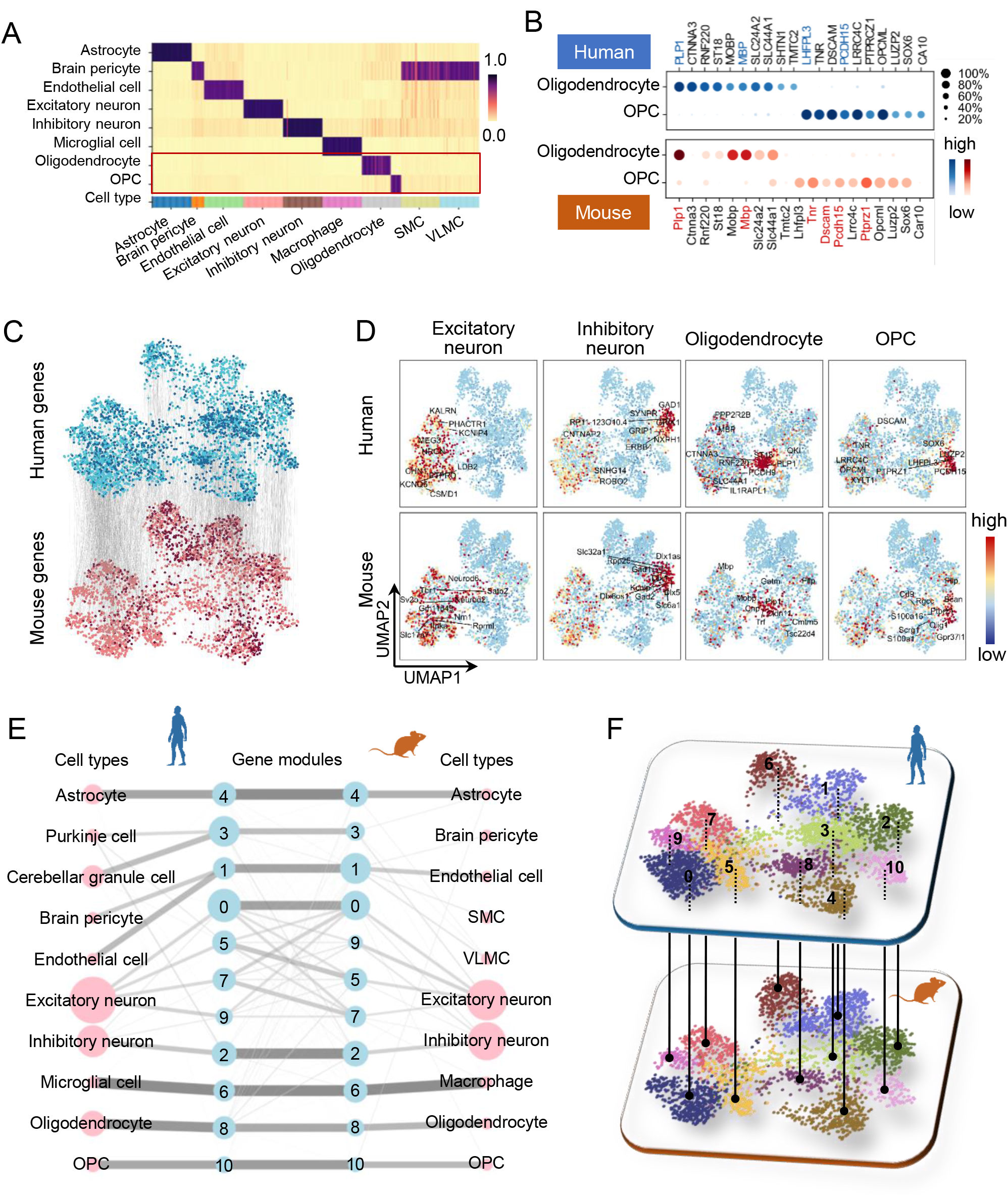
Application of CAME to human and mouse scRNA-seq data of brain cells. (**A**) The predicted cell-type probabilities for each cell (each column) in the mouse brain scRNA-seq data. The gene expressions of the human brain were taken as the reference. Each row indicates a cell type in human data. OPC is short for “oligodendrocyte precursor cells”, SMC is short for “smooth muscle cell”, and VLMC is short for “vascular and leptomeningeal cell”. (**B**) The top homologous DEG expressions of oligodendrocytes and (predicted) OPCs in human and mouse data, including several marker genes reported by previous literature (collected from CellMarker, colored by red or blue). (**C**) Cross-species alignment of the gene embeddings output by CAME, where each dot represents a gene and each edge indicates the homology between a pair of genes. Genes shared between species are colored by light-blue (human) or pink (mouse) while the other genes are colored by dark-blue (human) or dark-red (mouse). (**D**) The UMAP plots of gene embeddings showing the average expression patterns (z-scored across cell-types for each gene) of four cell types (excitatory neurons, inhibitory neurons, oligodendrocytes, OPCs) of human and mouse brains, where the color of each dot is scaled by the expression level of that cell type in the gene. (**E**) Abstracted graph of the heterogenous cell-gene graph, each node represents a cell type (pink) or a gene module (light blue). The size of a node is scaled by the number of single cells in that type or the number of genes in that gene module. The width of an edge is scaled by either the normalized mean expression levels of a cell type in the connected gene module or the conservancy of inter-species gene modules based on the gene embeddings learned by CAME. (**F**) Gene modules detected by joint module extraction of genes from humans (above) and mice (below).

Similar results were found when comparing four subtypes of the inhibitory neurons (VIP+, SST+, LAMP5+, PVALB+) between humans and mice. CAME still achieved a cell-typing accuracy of 98.3% and 95.5% for human-to-mouse and mouse-to-human label transfers, respectively, which are consistently higher than that of the benchmarking methods (93.4% and 92.0% for SciBet, 84.3% and 51.3% for SingleCellNet, 98.0% and 78.9% for Cell BLAST, 97.3% and 87.3% for ItClust, 69.5% and 78.9% for Scmap, 94.2% and 87.2% for Seurat) (**Supplementary Figure S12A and B**), although deferentially expressed genes (DEGs) for each homologous subtype seems not transferrable across species (**Supplementary Figure S12C and D**). The UMAP plots of cell embeddings showed that these major homologous cell-types were well aligned with each other. This suggested that the major types of brain cells in humans and mice are well conserved (**Supplementary Figure S9)**.

CAME also gave interpretable gene embeddings and enabled us to explore both intra- and inter-species relationships between genes. The UMAP plots of gene embeddings showed that the relative positions of human and mouse homologous genes were very consistent (**Figure 5C**). We further demonstrated the averaged gene expression profile on the UMAP plots of gene embeddings, where each point represents a gene (**Figure 4C** and **Supplementary Figure S11B**). It is worth noting that the neighbor genes tend to be co-expressed in the same cell types, such as those in excitatory, inhibitory neurons, oligodendrocytes, and OPCs (**Figure 4D**). There were more cell type-specific genes in human oligodendrocytes than in mice, indicating the evolutionary divergence between humans and mice. A population of genes was only detected in the human dataset, and most of them were associated with Purkinje cells and cerebellum granule cells, which were not detected in the mouse dataset due to their sources from different brain regions. These genes were arranged where there were few mouse genes around (**Figure 4F**, and **Supplementary Figure S11B**).

**Figure 5.**
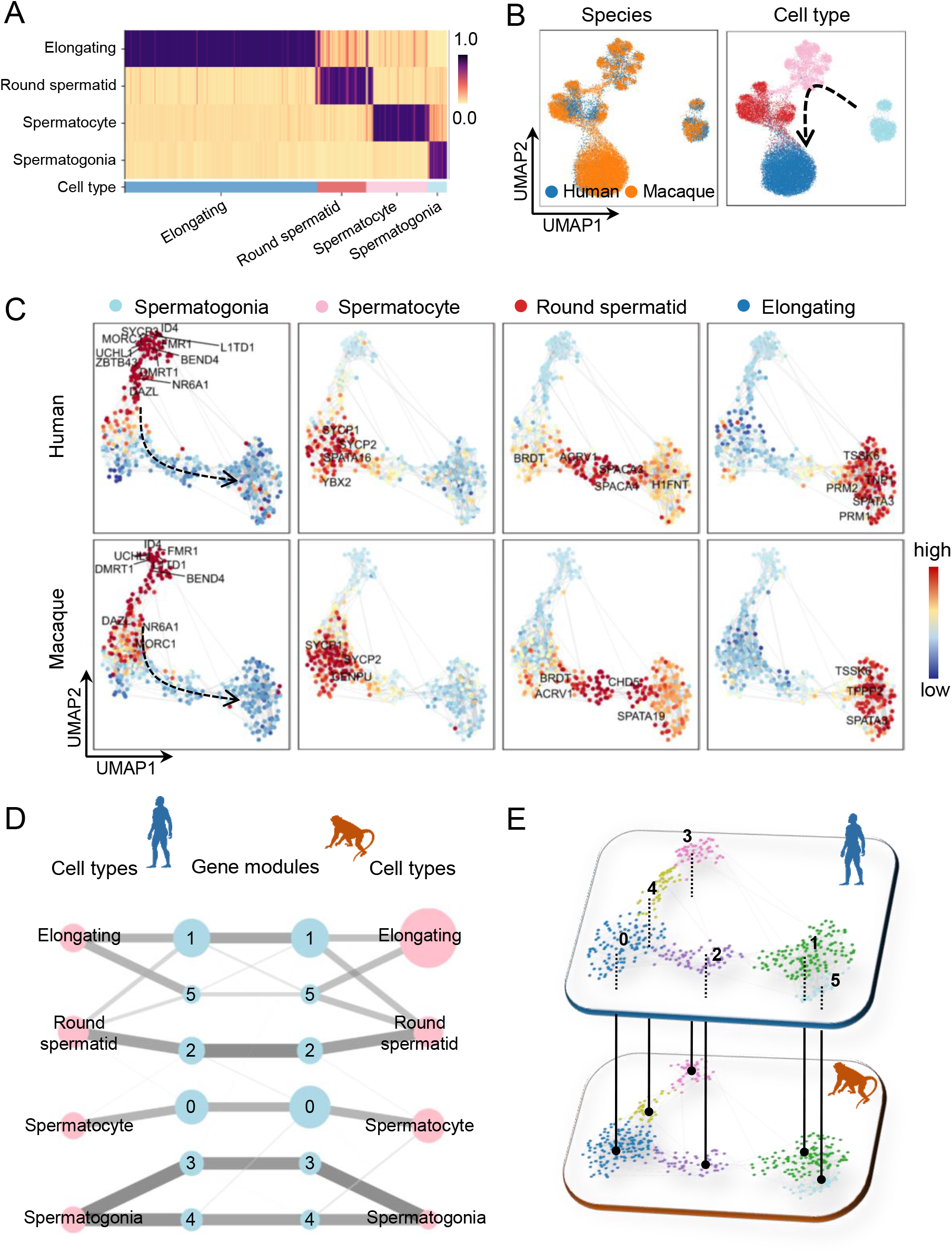
Application of CAME to human and macaque scRNA-seq data during spermatogenesis. (**A**) The predicted cell-type probabilities for each macaque testicular cell (each column). The gene expressions of human testis were taken as the reference. Each row indicates a cell type in the human data. (**B**) The UMAP plots of cell embeddings output by CAME, colored by datasets (left) or cell type (right). (**C**) 2D visualization of gene embeddings showing the average expression patterns (z-scored across cell-types for each gene) of the four stages across spermatogenesis, where each point represents a gene and the color of each scatter is scaled by the expression level of that cell type in the gene. (**D**) Abstracted graph of the heterogenous cell-gene graph. Each node represents a cell type (pink) or a gene module (light blue). The size of a node is scaled by the number of single cells in that type or the number of genes in that gene module. The width of an edge is scaled by either the normalized mean expression levels of a cell type in the connected gene module or the conservancy of inter-species gene modules based on the gene embeddings learned by CAME. (**E**) Gene modules detected by joint module extraction of genes from humans (above) and macaques (below).

The aligned gene embeddings across species can facilitate us to jointly extract cell type-specific gene modules with different degrees of conservancies between species, and each module corresponds to a cell type like OPCs, or related cell types like endothelial cells and its subtypes (**Figure 4E** and **Methods**). As expected, based on gene ontology (GO) [33, 34] enrichment analysis, we found that the functions associated with most homologous gene modules were generally consistent with each other (**Supplementary Table S2**). For example, both the human and mouse genes in module 2 (which was associated with inhibitory neurons) tended to relate functions like “forebrain neuron differentiation” and “learning or memory”. Both the human and mouse genes in module 6 (corresponding human microglia and mouse macrophage) were related to functions like “positive regulation of cytokine production”, and “leukocyte migration”. By contrast, the function “ventral spinal cord development” was only enriched in human module 3 but not in mice, considering their gene members were quite different; though they were both associated with the function “cell differentiation in hindbrain” and “cerebellar cortex formation”.

### CAME could reveal conservative expression dynamics during spermatogenesis between human and macaque

Comparison of continuous biological processes between two species is of much interest in evolutionary biology. We applied CAME to two scRNA-seq datasets from human and macaque testicular single cells [9] with the former as the reference one. CAME achieved a very distinct cell-typing accuracy of 95.0% (86.0% for SciBet, 89.2% for SingleCellNet, 76.1% for Cell BLAST, 53.4% for ItClust, 87.3% for Scmap, 89.1% for Seurat), and a precise alignment of the homologous cell types of human and macaque with each other (**Figure 5A and B**). Besides, the labeled spermatogonia, spermatocyte, round spermatid, and elongating cells are correctly merged along the underlying differentiation trajectory. This suggested that CAME could well decipher the conserved four-stage spermatogenesis processes of humans and macaques.

Very interestingly, the continuously dynamic changing process of spermatogenesis can also be revealed by the UMAP plot of gene embeddings (**Figure 5C**). As illustrated, CAME extracted four sets of genes, including some typical marker ones [32], that are highly co-expressed in the four main stages of spermatogenesis and form well-organized expression dynamics, suggesting the order of critical gene activations during spermatogenesis (**Figure 5C**). By joint extraction of gene modules, we found that the four stages of spermatogenesis were quite conservative from the aspect of gene modules (**Figure 5D and E**). For example, modules 3, 4, and 0 were highly expressed in spermatogonia and spermatocyte respectively for both humans and macaques. And round spermatids and elongating spermatids shared modules 2, 1, and 5 in different degrees. Typically, both human and macaque module 4 was associated with functions like “RNA splicing”, and module 1 was associated with “sperm motility” and “spermatid development/differentiation”, which were typical characteristics of elongating spermatids (**Supplementary Table S3**).

## Conclusions

Cross-species comparative and integrative analysis at single-cell resolution has deepened our understanding of the origin and evolutionary mechanisms of cellular states. Exploring the conservative and divergent characteristics of homologous cell states between human and other model and non-model species, for example, can help us to determine the animal model for studying human disease [5-7].

However, in addition to technical noises, the systematic shift of gene expressions associated with distinct species and the uncertainty of the orthologous genes make it much more difficult than within-species data integration. Moreover, existing approaches for cross-species integration were mainly based on one-to-one homologous genes. However, when it is needed to align cell types across long distant species, especially when a large number of gene duplications were involved during the evolution process [27,28], considering only the one-to-one homologous genes will inevitably lose a lot of important information. Even so, cells of homologous types are thought to have similar expression patterns, that is, they may co-express a cell type-specific combination of genes. These genes may not be easy to be identified as the marker genes with high expression levels but can act as “bridges” between cells that co-expressed them. Besides, the gene-homology mappings can bridge the gene nodes of two species, where the non-one-to-one homologies can also be used.

Thus, we take the gene expression matrix as a bipartite graph with cell and gene nodes and utilize the gene homologies to form a multipartite graph. Based on this, we proposed CAME to utilize a cell-gene heterogeneous graph neural network to facilitate the “message-passing” from one species to the other. CAME can achieve the alignment of both cells and genes from different species. As a result, CAME can not only achieve accurate and robust cell-type assignment, but also reveal biological insights into the conservative and divergent characteristics between species. When handling multiple references, most integration approaches have to perform pairwise alignment for individual batches, where the order of pairwise alignment can affect the results and the computational complexity rises quadratically with the number of batches. Others like Harmony [30] and Cell BLAST [17] are capable to align multiple datasets simultaneously. We demonstrated that CAME can remove batch effects for multiple references even when batch labels are not provided. This is an important characteristic for integrating various datasets and constructing a unified cell-typing reference.

It should be noticed that the heterogeneous graph neural network structure of CAME can also be applied to the scenario of within-species data integration, or when we consider only the one-to-one homologous genes. The only adjustment is to replace each gene-gene edge with a single gene node. Moreover, this strategy can be applied for multi-omics label transfer and data integration. In summary, we believe that CAME will serve as a powerful tool for integrative and comparative analysis across species as well as multi-omics integration.

## Methods

### Build a heterogeneous cell-gene graph

Let’s denote a gene expression matrix with *N* cells and *M* genes as *X = (X*_1_, *X*_2_, …, *X*_*N*_)^*T*^ *∈ ℝ*^*N×M*^, where each row *X*_*i*_ *= (x*_*i*1_, *x*_*i*2_, … *x*_*iM*_) *∈ ℝ*^*M*^ with an element *x*_*ij*_ representing the (normalized) expression value of a cell *i* in a gene *j*. We take 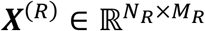 and 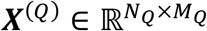 as the reference and query datasets respectively, 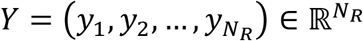 as the cell-type labels of the reference dataset and a set of gene pairs {(*g*_*i*_, *g*_*j*_)}_*ij*_ to indicate the homology between two species. Note that *M*_*R*_ is not necessarily equal to *M*_*Q*_.

The reference and query expression matrices and the homology together are represented as a heterogeneous cell-gene graph with each node acting as a cell or a gene (**Figure 1A**). A cell-gene edge in the graph indicates that this cell has non-zero expression of the gene, a gene-gene edge indicates a homology between each other, and a cell-cell edge indicates the expression profiles of these two cells are similar to each other. In other words, in this graph, there are two types of nodes, cell and gene, and six types of edges (relations) including “a cell expresses a gene”, “a gene is expressed by a cell”, “cell-cell similarity”, “gene-gene homology”, “cell self-loop” and “gene self-loop”.

### Design a heterogeneous graph neural network

CAME adopts a heterogeneous graph neural network, which was motivated by a relational graph convolutional network [24] for a graph of homogeneous nodes but heterogeneous edges. We denote the convolution weights for these six edge types as *W*_*cg*_, *W*_*gc*_, *W*_*cc*_, *W*_*gg*_, *W*_*c*_ and *W*_*g*_, respectively (**Figure 1B**). For each cell *i*, its initial embedding (the 0-th layer) is calculated as:

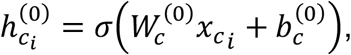

where *σ* is the ReLU activation function; 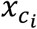 is the gene expressions in the cell *i* (one-to-one homologous genes are taken as the common input features) and 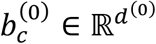 is the learnable bias vector. The genes, however, lack the initial embeddings in the 0-th layer and can be aggregated from their neighbor cells as follows:

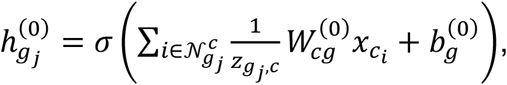

where 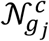 is the set of cells that have expressed the gene *j*, and 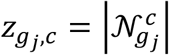 is the normalization factor. This approach keeps the number of model parameters stay constant to the number of genes, which differs from the commonly used initialization that assigns a learnable embedding for those nodes without input features, where the increasing number of model parameters might lead to an overfitted model. It can also allow inductive learning for the genes not involved in the training process.

While in each hidden layer *l ≥* 1, the node features for the cell *i* and the gene *j* can be calculated as:

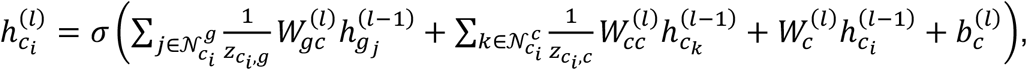

and

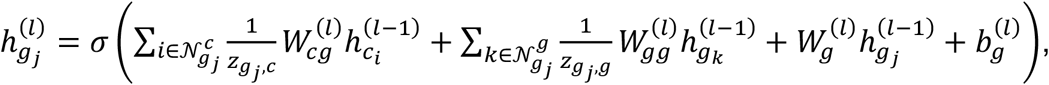

respectively. Note that we treat the edges between homologous genes and the self-loop on each gene identically, i.e., 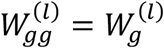. To boost the ‘message’ flow between reference and query nodes, we adopt a recurrent convolution, where the parameters are shared across the hidden layers, that is, 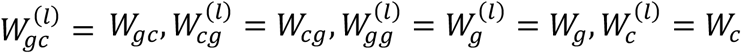 and 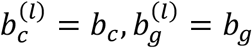 for 1 ≤ *l* ≤ *L*, where *L* is the total number of the hidden layers. We recommend to set *L* as 2 or 3 in practice, and the default setting is 2. We also adopt the layer normalization for all the hidden states to facilitate fast training convergence and high performance (**Supplementary Figure S1**).

When it comes to the cell-type classifier, we adopt the attention mechanism for graph convolution [25], where each cell pays distinct attention to its neighbor genes. Specifically, for each cell *i*, the output states 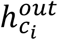 for cell-type identification is aggregated from their neighbor genes:

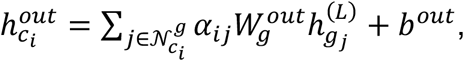

where *α*_*ij*_ is the attention that the cell *i* pays to the gene *j*, calculated as:

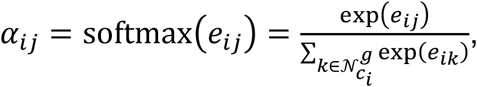

with

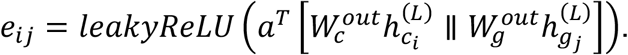

In addition, we use multi-head attention to enhance the model capacity and robustness, where there are several attention-heads with their own parameters, and their outputs are merged by taking averages:

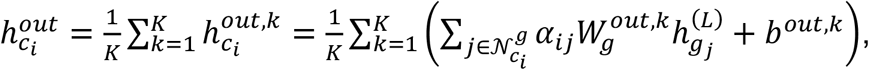

where *K* is the total number of attention-heads, set as 8 by default.

Finally, the output layer states for cell-type classification were normalized in two different ways: (1) the *softmax* function over cell types for multi-class classification:

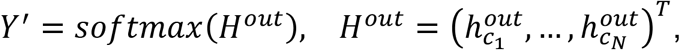

where *Y*′ *∈ ℝ*^*N*×*T*^ and each row is the predicted probabilities over the *T* cell types for a cell; (2) the sigmoid function for multi-label classification:

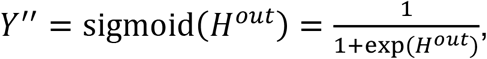

where *Y*′′ *∈ ℝ*^*N*×*T*^ and each element 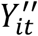 is the predicted probability of the cell type *t* for the cell *i*.

### The classification loss and label smoothing

The classification loss for cells in reference datasets is calculated by the weighted cross-entropy loss combined with *L*_2_ regularization as below:

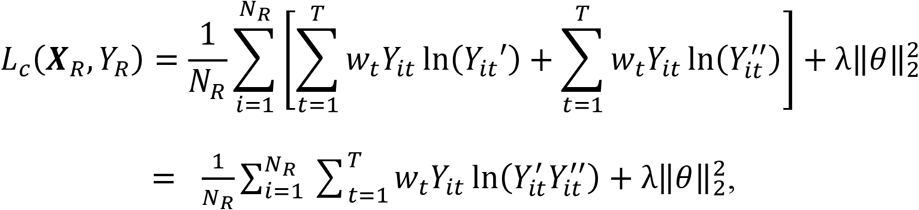

where *w*_*t*_ is the class-weight for cell-type *t* satisfying 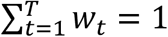 To avoid the model being dominated by the major populations and ignoring those rare types, we set 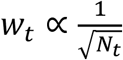 and *N*_*t*_ is the number of cells of cell type *t* in the reference dataset. *θ* represents all the learnable parameters and *λ*_1_ is the penalization coefficient that controls the power of *L*_2_ regularization, and the default value of *λ*_1_ is 0.01.

To prevent the model from being overconfident and improve the stability and generalization of the model, we utilize label smoothing [35]. We minimize the cross-entropy between the modified targets 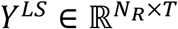 and the model outputs *Y*′, where 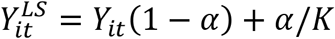, and the final objective function is as below:

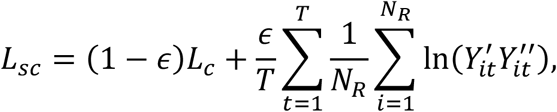

where *∈* controls the degree of smoothness, set as 0.1 by default. Finally, CAME adopts Adam optimizer [36] with a learning rate of 0.001 for training.

### Checkpoint selection

When training the heterogeneous graph neural network, we would like to choose the epoch where the classification result of query datasets achieves the highest accuracy. However, in practice, the exact type labels of the query cells are unknown, hindering us from choosing the best model. We put forward a metric to approximate the accuracy. Specifically, we first cluster the query cells to get the pseudo-labels *Y*^*cluster*^ for the query cells and introduce adjusted mutual information (AMI) [37] to account for the chance between the model-predicted cell-type labels and the pseudo-labels of the query cells to help decide when to stop. AMI is defined as

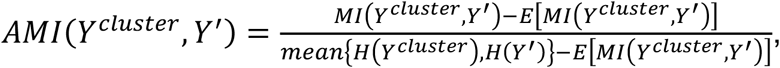

where *H(X*) is the entropy of *X, MI(X, Y*) is the mutual information between variables *X* and *Y*. *E[MI(Y*^*cluster*^, *Y*′)] is the expected mutual information based on a “permutation model” [38], in which cluster labels are generated randomly subject to having a fixed number of clusters and points in each cluster. We think that a well-trained model is expected to preserve the intrinsic data structure so that the predicted labels should be highly consistent with the pseudo-labels to some extent. We run the model with 400 epochs and choose the checkpoint with the largest AMI. The clustering process will be described in the section “*pre-clustering of the query cells*” in detail.

### Training using the mini-batches on sub-graphs

When training CAME on the graphics processing unit (GPU), the size of a dataset will be limited by the GPU memory. For example, training CAME on 100,000 cells could take about 13.75GB of memory, which exceeds the graphic memory of most GPUs. To handle this issue, we utilized a mini-batch training process by using the graph segmentation technique. Specifically, we first randomly divided all the cells (including cells in reference and query) into several groups, taken as mini-batches. For each mini-batch, we created a node-induced subgraph for a given group of cells, which contains all the cells in this group and all the genes expressed by these cells. Then, we iterated all subgraphs and feed the subgraphs to the graph neural network one by one. All the parameters will be updated for each mini-batch training process. We performed extensive experiments by using mini-batch training process and found it is suitable to choose batch-size as 8192 or more, for that it achieved the comparable accuracy compared with the whole graph training (**Supplementary Figure S14A**) and the cost of GPU memory stays constant (2.4GB) for datasets at different scales (**Supplementary Figure S14B**). Such low consumption of graphic memory means you can use CAME on almost all graphics cards. It is worth noting that the runtime of the batch-training process will be largely increased (**Supplementary Figure S14B**) since we cannot feed forward the whole graph on a single epoch.

### Preprocessing of the single-cell datasets

For each scRNA-seq dataset, we first normalized the counts of each cell by its library size with a scale factor multiplied (10,000 by default) and log-transformed with a pseudo-count added for the downstream analysis.

### Gene selection

Highly variable genes (HVGs) and deferentially expressed genes (DEGs) are generally thought to be highly informative and the latter is especially useful for cell-type characterization. Therefore, we used both HVGs and DEGs and extended them using homologous mappings to form the highly informative gene (HIG) sets for constructing the heterogeneous graph. We adopted the same approach as used in Seurat-v2 [39] with ScanPy [40] built-in function highly_variable_genes() to identify HVGs, separately from both reference and query data. Specifically speaking, it calculated the average expression and dispersion (variance/mean) for each gene and placed these genes into several bins based on the (log-transformed) average expression. The normalized dispersions were then obtained by scaling with the mean and standard deviation of the dispersions within each bin. We selected the top 2000 genes with the highest dispersions as HVGs of that dataset. We computed the DEGs separately for reference and query dataset by Student’s t-test, which is done through rank_genes_groups() function from the ScanPy package [40]. For reference data, cells are grouped by their cell-type labels, while for the query data, cells are grouped by their pseudo-labels, i.e., the pre-clustering labels.

Genes used as the cell-node features should be shared between species (or datasets). For both reference and query datasets, we first took the top 50 DEGs for each cell group and retained genes with one-to-one homology in the other species. We then took the union of the resulting two sets of genes for input. The resulting number of genes used for defining cell-node features ranges from 240 to 400 for distant species pairs (human to zebrafish for example) and from 400 to 900 for the others.

We combined both HVGs and DEGs from reference and query data to decide the node genes used for training the graph neural network. Specifically, we first took the union of the HVGs and DEGs for each dataset, denoted as *g*_*r*_ and *g*_*q*_ for reference and query respectively. Then we extracted the genes that have homologies in *g*_*r*_ from the query data, and the homologous genes for *g*_*q*_ from the reference data denoted as 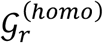 and 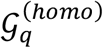 respectively. Finally, we determined 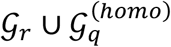, the union of *g*_*r*_ and 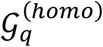, as the node genes for the reference species and 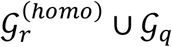 as the node genes for the query species. The tables containing gene homology information for each species pair were downloaded from the BioMart web server (http://www.ensembl.org/biomart/martview) [41].

### Construction of the single-cell graphs based on KNNs

The normalized expression matrices were centralized and scaled within each dataset, followed by principal component analysis (PCA) to reduce the dimensionality. We searched approximate KNNs for each cell based on the top 30 PCs with the highest explained variances. We adopted *k = 5* neighbors for each cell to make the graph sparse enough for computational efficiency. These neighbor connections provided “cell-cell” edges as a part of the heterogeneous graph.

### Pre-clustering of the query cells

To facilitate model selection, we pre-clustered the query cells using a graph-based clustering method, that is, performing community detection using the Leiden algorithm [42] on the single-cell KNN graph. We constructed the KNN graph in almost the same way as described above, except that the number of neighbors *k* was set as 20 and the clustering resolution is set as 0.4 by default.

### Unifying cell-type labels across datasets

For data downloaded from the Cell BLAST web server [17], the cell-type labels were already unified by Cell Ontology [43], a structured vocabulary for cell types. While for unifying annotations from the other datasets, we directly referred to Cell Ontology and manually adjusted the annotations. The annotations were used as ground truth.

### Gene module extraction

To extract cell type-specific gene modules shared between species, we took all the gene embeddings (of both species) on the last hidden layer and performed KNN searching for each gene. Like clustering cells, we performed Leiden community detection on the KNN graph of genes. The clustering resolution was set as 0.8 by default.

### Calculating weights between gene modules

The weights *S*_*ij*_ between homologous gene modules *Mod*_*i*_ and *Mod*_*j*_ on the abstracted graph were calculated as follows:

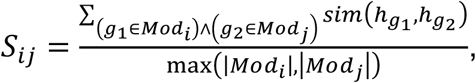

where *h*_*g*_ is the embedding vector of gene *g* and *sim(⋅,⋅*) is the similarity function, cosine similarity by default. |*Mod*| represents the number of genes in this module.

### Benchmarking cell-type assignment

For benchmarking cell-type assignment, we collected 54 scRNA-seq datasets from five tissues across seven different species (**Supplementary Figure 3A, Supplementary Table 1**), paired datasets of different species within the same tissue, and filtered those pairs where more than 50% of query cells are unresolved in the reference cell types, resulting 649 cross-species dataset pairs. For each dataset, we removed the cell types of less than 10 cells. CAME was compared with six benchmarking methods including Seurat V3 [16], ItClust [18], Scmap [44], SingleCellNet [14], SciBet [15], and Cell BLAST [17]. For Seurat V3, we input the raw data, used the default normalize process by NormalizeData() function, extracted the top 2000 HVGs by its FindVariableFeatures() function for reference and query respectively, and performed further annotation process as described in its documentation. For ItClust, since it provides an automatic workflow including preprocessing and annotation, we input the raw data. For Scmap, we log-transformed the raw counts with pseudo-count 1 added and used its inherited function selectFeaures() to select the top 2000 HVGs with a threshold=0.1 in function scmapCluster() (which works better for the cross-species scenario than its default value). For SingleCellNet, we also input raw data as it suggested, used splitCommon function to split for training and assessment, employed expTMraw function to transform training data, and then used scn_predict to make predictions for query dataset. For SciBet, we used R to perform all the operations. We first input the library-size-normalized data calculated by cpm() function of package edgeR [45] and used SelectGene_R() function from SciBet package to select 2000 HVGs, and used SciBet_R() function to annotate the query data. For Cell BLAST, we used the raw data as input and used find_variable_genes() to select HVGs with default parameters and took the union of the HVGs between reference and query datasets. After that, the datasets were combined together to remove their batch effects by using function fit_DIRECTi() with lambad_reg=0.001 as suggested by the original authors to stabilize the training process. Cell BLAST also provides a supervised training process that leverages the cell type labels of reference datasets to perform label transfer. However, it led to a 4% decrease in the average accuracy compared with their previous batch effects correction process. All hyper-parameters not mentioned were set with default values in these six packages.

To evaluate the performance of the cell-type assignment, we adopted three metrics: Accuracy, MarcoF1, and WeightedF1. Accuracy is the most common criterion and it directly measures how many of the predictions are the same as the actual ones:

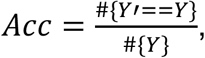

where *#* is the sign of cardinality. Specifically, *#{Y*_*true*_} means the number of the total cells and *#{Y*′ == *Y*} means the number of correctly predicted ones.

We also used *MacroF*1 and *Weightee*1 which consider the *F*_1_-score for each cell type. For a binary classification task, precision and recall are calculated as

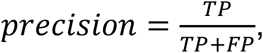

and

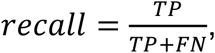

respectively, where TP, FP, and FN represent the number of true positives, false positives, and false negatives, respectively.

The *F*_1_-score is the harmonic mean of *precision* and *recall*:

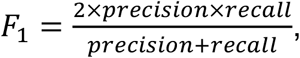

and the *MacroF*1 is defined as the average of class-wise *F*_1_-scores,

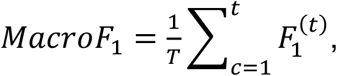

where 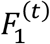 represents the *F*_1_ -score for cell type *t*. The *WeightedF*1 considers the proportion of each class,

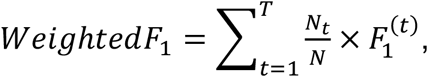

where *N*_*t*_*/N* represents the proportion of type *t* in all cells.

### Benchmarking data integration

FastMNN [46], Harmony [30], and Seurat-v3 [16] were performed using the corresponding R package through SeuratWrapper, following the online documents with default settings. FastMNN, Harmony, and Seurat shared the same normalization and the top 2000 HVGs by Seurat function NormalizeData and FindVariableFeatures, respectively. Harmony was performed on the PCA-reduced embeddings. The number of reduced dimensions for these three methods was set as 50 for all pairs of datasets. LIGER [19] took the raw count data as input and run with the default pipeline. Cell BLAST [17] was performed using its Python package, following the standard pipeline with the default settings.

## Acknowledgments

This work has been supported by the National Key Research and Development Program of China [2019YFA0709501]; the Strategic Priority Research Program of the Chinese Academy of Sciences (CAS) [XDPB17], the Key-Area Research and Development of Guangdong Province [2020B1111190001], the National Natural Science Foundation of China [61621003]; the National Ten Thousand Talent Program for Young Top-notch Talents, the CAS Frontier Science Research Key Project for Top Young Scientist [QYZDB-SSW-SYS008], and the Shanghai Municipal Science and Technology Major Project [2017SHZDZX01].

## References

[1] E.Z. Macosko, A. Basu, R. Satija, J. Nemesh, K. Shekhar, M. Goldman, I. Tirosh, A.R. Bialas, N. Kamitaki, E.M. Martersteck, J.J. Trombetta, D.A. Weitz, J.R. Sanes, A.K. Shalek, A. Regev, S.A. McCarroll, Highly Parallel Genome-wide Expression Profiling of Individual Cells Using Nanoliter Droplets, Cell 161(5) (2015) 1202–1214.

[2] A.A. Kolodziejczyk, J.K. Kim, V. Svensson, J.C. Marioni, S.A. Teichmann, The technology and biology of single-cell RNA sequencing, Molecular cell 58(4) (2015) 610–20.

[3] J.C. Marioni, D. Arendt, How Single-Cell Genomics Is Changing Evolutionary and Developmental Biology, Annual Review of Cell and Developmental Biology 33(1) (2017) 537–553.

[4] M.E.R. Shafer, Cross-Species Analysis of Single-Cell Transcriptomic Data, Front Cell Dev Biol 7 (2019) 175–175.

[5] E. Drokhlyansky, C.S. Smillie, N. Van Wittenberghe, M. Ericsson, G.K. Griffin, G. Eraslan, D. Dionne, M.S. Cuoco, M.N. Goder-Reiser, T. Sharova, O. Kuksenko, A.J. Aguirre, G.M. Boland, D. Graham, O. Rozenblatt-Rosen, R.J. Xavier, A. Regev, The Human and Mouse Enteric Nervous System at Single-Cell Resolution, Cell 182(6) (2020) 1606-1622.e23.

[6] L. Geirsdottir, E. David, H. Keren-Shaul, A. Weiner, S.C. Bohlen, J. Neuber, A. Balic, A. Giladi, F. Sheban, C.-A. Dutertre, C. Pfeifle, F. Peri, A. Raffo-Romero, J. Vizioli, K. Matiasek, C. Scheiwe, S. Meckel, K. Mätz-Rensing, F. van der Meer, F.R. Thormodsson, C. Stadelmann, N. Zilkha, T. Kimchi, F. Ginhoux, I. Ulitsky, D. Erny, I. Amit, M. Prinz, Cross-Species Single-Cell Analysis Reveals Divergence of the Primate Microglia Program, Cell 179(7) (2019) 1609-1622.e16.

[7] R.D. Hodge, T.E. Bakken, J.A. Miller, K.A. Smith, E.R. Barkan, L.T. Graybuck, J.L. Close, B. Long, N. Johansen, O. Penn, Z. Yao, J. Eggermont, T. Höllt, B.P. Levi, S.I. Shehata, B. Aevermann, A. Beller, D. Bertagnolli, K. Brouner, T. Casper, C. Cobbs, R. Dalley, N. Dee, S.L. Ding, R.G. Ellenbogen, O. Fong, E. Garren, J. Goldy, R.P. Gwinn, D. Hirschstein, C.D. Keene, M. Keshk, A.L. Ko, K. Lathia, A. Mahfouz, Z. Maltzer, M. McGraw, T.N. Nguyen, J. Nyhus, J.G. Ojemann, A. Oldre, S. Parry, S. Reynolds, C. Rimorin, N.V. Shapovalova, S. Somasundaram, A. Szafer, E.R. Thomsen, M. Tieu, G. Quon, R.H. Scheuermann, R. Yuste, S.M. Sunkin, B. Lelieveldt, D. Feng, L. Ng, A. Bernard, M. Hawrylycz, J.W. Phillips, B. Tasic, H. Zeng, A.R. Jones, C. Koch, E.S. Lein, Conserved cell types with divergent features in human versus mouse cortex, Nature 573(7772) (2019) 61–68.

[8] A. Sebé-Pedrós, E. Chomsky, K. Pang, D. Lara-Astiaso, F. Gaiti, Z. Mukamel, I. Amit, A. Hejnol, B.M. Degnan, A. Tanay, Early metazoan cell type diversity and the evolution of multicellular gene regulation, Nat Ecol Evol 2(7) (2018) 1176–1188.

[9] A.N. Shami, X. Zheng, S.K. Munyoki, Q. Ma, G.L. Manske, C.D. Green, M. Sukhwani, K.E. Orwig, J.Z. Li, S.S. Hammoud, Single-Cell RNA Sequencing of Human, Macaque, and Mouse Testes Uncovers Conserved and Divergent Features of Mammalian Spermatogenesis, Developmental Cell (2020).

[10] M.A. Tosches, T.M. Yamawaki, R.K. Naumann, A.A. Jacobi, G. Tushev, G. Laurent, Evolution of pallium, hippocampus, and cortical cell types revealed by single-cell transcriptomics in reptiles, Science (New York, N.Y.) 360(6391) (2018) 881–888.

[11] J. Wang, H. Sun, M. Jiang, J. Li, P. Zhang, H. Chen, Y. Mei, L. Fei, S. Lai, X. Han, X. Song, S. Xu, M. Chen, H. Ouyang, D. Zhang, G.-C. Yuan, G. Guo, Tracing cell-type evolution by cross-species comparison of cell atlases, Cell Reports 34(9) (2021) 108803.

[12] A.W. Zhang, C. O’Flanagan, E.A. Chavez, J.L.P. Lim, N. Ceglia, A. McPherson, M. Wiens, P. Walters, T. Chan, B. Hewitson, D. Lai, A. Mottok, C. Sarkozy, L. Chong, T. Aoki, X. Wang, A.P. Weng, J.N. McAlpine, S. Aparicio, C. Steidl, K.R. Campbell, S.P. Shah, Probabilistic cell-type assignment of single-cell RNA-seq for tumor microenvironment profiling, Nature Methods 16(10) (2019) 1007–1015.

[13] X. Shao, J. Liao, X. Lu, R. Xue, N. Ai, X. Fan, scCATCH: Automatic Annotation on Cell Types of Clusters from Single-Cell RNA Sequencing Data, iScience 23(3) (2020) 100882.

[14] Y. Tan, P. Cahan, SingleCellNet: A Computational Tool to Classify Single Cell RNA-Seq Data Across Platforms and Across Species, Cell Systems 9(2) (2019) 207-213.e2.

[15] C. Li, B. Liu, B. Kang, Z. Liu, Y. Liu, C. Chen, X. Ren, Z. Zhang, SciBet as a portable and fast single cell type identifier, Nature Communications 11(1) (2020) 1818.

[16] T. Stuart, A. Butler, P. Hoffman, C. Hafemeister, E. Papalexi, W.M. Mauck, 3rd, Y. Hao, M. Stoeckius, P. Smibert, R. Satija, Comprehensive Integration of Single-Cell Data, Cell 177(7) (2019) 1888-1902.e21.

[17] Z.-J. Cao, L. Wei, S. Lu, D.-C. Yang, G. Gao, Searching large-scale scRNA-seq databases via unbiased cell embedding with Cell BLAST, Nature Communications 11(1) (2020) 3458.

[18] J. Hu, X. Li, G. Hu, Y. Lyu, K. Susztak, M. Li, Iterative transfer learning with neural network for clustering and cell type classification in single-cell RNA-seq analysis, Nature Machine Intelligence 2(10) (2020) 607–618.

[19] J.D. Welch, V. Kozareva, A. Ferreira, C. Vanderburg, C. Martin, E.Z. Macosko, Single-Cell Multi-omic Integration Compares and Contrasts Features of Brain Cell Identity, Cell 177(7) (2019) 1873-1887.e17.

[20] L. Zhang, S. Zhang, Learning common and specific patterns from data of multiple interrelated biological scenarios with matrix factorization, Nucleic acids research 47(13) (2019) 6606–6617.

[21] D. Arendt, J.M. Musser, C.V.H. Baker, A. Bergman, C. Cepko, D.H. Erwin, M. Pavlicev, G. Schlosser, S. Widder, M.D. Laubichler, G.P. Wagner, The origin and evolution of cell types, Nature reviews. Genetics 17(12) (2016) 744–757.

[22] S. Aibar, C.B. González-Blas, T. Moerman, V.A. Huynh-Thu, H. Imrichova, G. Hulselmans, F. Rambow, J.C. Marine, P. Geurts, J. Aerts, J. van den Oord, Z.K. Atak, J. Wouters, S. Aerts, SCENIC: single-cell regulatory network inference and clustering, Nat Methods 14(11) (2017) 1083–1086.

[23] M.C. Oldham, S. Horvath, D.H. Geschwind, Conservation and evolution of gene coexpression networks in human and chimpanzee brains, Proceedings of the National Academy of Sciences of the United States of America 103(47) (2006) 17973–8.

[24] M. Schlichtkrull, T.N. Kipf, P. Bloem, R. van den Berg, I. Titov, M. Welling, Modeling Relational Data with Graph Convolutional Networks, in: A. Gangemi, R. Navigli, M.-E. Vidal, P. Hitzler, R. Troncy, L. Hollink, A. Tordai, M. Alam (Eds.) The Semantic Web, Springer International Publishing, Cham, 2018, pp. 593–607.

[25] P. Veličković, G. Cucurull, A. Casanova, A. Romero, P. Liò, Y.J.a.e.-p. Bengio, Graph Attention Networks, 2017, p. 1710.10903.

[26] T. Hoang, J. Wang, P. Boyd, F. Wang, C. Santiago, L. Jiang, S. Yoo, M. Lahne, L.J. Todd, M. Jia, C. Saez, C. Keuthan, I. Palazzo, N. Squires, W.A. Campbell, F. Rajaii, T. Parayil, V. Trinh, D.W. Kim, G. Wang, L.J. Campbell, J. Ash, A.J. Fischer, D.R. Hyde, J. Qian, S. Blackshaw, Gene regulatory networks controlling vertebrate retinal regeneration, Science (New York, N.Y.) 370(6519) (2020).

[27] V. Ravi, B. Venkatesh, The Divergent Genomes of Teleosts, Annual review of animal biosciences 6 (2018) 47–68.

[28] S.M. Glasauer, S.C. Neuhauss, Whole-genome duplication in teleost fishes and its evolutionary consequences, Molecular genetics and genomics: MGG 289(6) (2014) 1045–60.

[29] B. Tasic, Z. Yao, L.T. Graybuck, K.A. Smith, T.N. Nguyen, D. Bertagnolli, J. Goldy, E. Garren, M.N. Economo, S. Viswanathan, O. Penn, T. Bakken, V. Menon, J. Miller, O. Fong, K.E. Hirokawa, K. Lathia, C. Rimorin, M. Tieu, R. Larsen, T. Casper, E. Barkan, M. Kroll, S. Parry, N.V. Shapovalova, D. Hirschstein, J. Pendergraft, H.A. Sullivan, T.K. Kim, A. Szafer, N. Dee, P. Groblewski, I. Wickersham, A. Cetin, J.A. Harris, B.P. Levi, S.M. Sunkin, L. Madisen, T.L. Daigle, L. Looger, A. Bernard, J. Phillips, E. Lein, M. Hawrylycz, K. Svoboda, A.R. Jones, C. Koch, H. Zeng, Shared and distinct transcriptomic cell types across neocortical areas, Nature 563(7729) (2018) 72–78.

[30] I. Korsunsky, N. Millard, J. Fan, K. Slowikowski, F. Zhang, K. Wei, Y. Baglaenko, M. Brenner, P.R. Loh, S. Raychaudhuri, Fast, sensitive and accurate integration of single-cell data with Harmony, Nat Methods 16(12) (2019) 1289–1296.

[31] L. McInnes, J. Healy, J.J.a.e.-p. Melville, UMAP: Uniform Manifold Approximation and Projection for Dimension Reduction, 2018, p. 1802.03426.

[32] X. Zhang, Y. Lan, J. Xu, F. Quan, E. Zhao, C. Deng, T. Luo, L. Xu, G. Liao, M. Yan, Y. Ping, F. Li, A. Shi, J. Bai, T. Zhao, X. Li, Y. Xiao, CellMarker: a manually curated resource of cell markers in human and mouse, Nucleic acids research 47(D1) (2019) D721–d728.

[33] The Gene Ontology resource: enriching a GOld mine, Nucleic acids research 49(D1) (2021) D325–d334.

[34] M. Ashburner, C.A. Ball, J.A. Blake, D. Botstein, H. Butler, J.M. Cherry, A.P. Davis, K. Dolinski, S.S. Dwight, J.T. Eppig, M.A. Harris, D.P. Hill, L. Issel-Tarver, A. Kasarskis, S. Lewis, J.C. Matese, J.E. Richardson, M. Ringwald, G.M. Rubin, G. Sherlock, Gene ontology: tool for the unification of biology. The Gene Ontology Consortium, Nature genetics 25(1) (2000) 25–9.

[35] C. Szegedy, V. Vanhoucke, S. Ioffe, J. Shlens, Z. Wojna, Rethinking the Inception Architecture for Computer Vision, 2016 IEEE Conference on Computer Vision and Pattern Recognition (CVPR), 2016, pp. 2818–2826.

[36] D.P. Kingma, J.J.C. Ba, Adam: A Method for Stochastic Optimization, abs/1412.6980 (2015).

[37] N.X. Vinh, J. Epps, J. Bailey, Information Theoretic Measures for Clusterings Comparison: Variants, Properties, Normalization and Correction for Chance, 11 (2010) 2837–2854.

[38] H. Ahrens, Lancaster, H. O.: The Chi-squared Distribution. Wiley & Sons, Inc., New York 1969. X, 366 S., 140 s, 13(5) (1971) 363–364.

[39] A. Butler, P. Hoffman, P. Smibert, E. Papalexi, R. Satija, Integrating single-cell transcriptomic data across different conditions, technologies, and species, Nature biotechnology 36(5) (2018) 411–420.

[40] F.A. Wolf, P. Angerer, F.J. Theis, SCANPY: large-scale single-cell gene expression data analysis, Genome Biol 19(1) (2018) 15.

[41] R.J. Kinsella, A. Kähäri, S. Haider, J. Zamora, G. Proctor, G. Spudich, J. Almeida-King, D. Staines, P. Derwent, A. Kerhornou, P. Kersey, P. Flicek, Ensembl BioMarts: a hub for data retrieval across taxonomic space, Database: the journal of biological databases and curation 2011 (2011) bar030.

[42] V.A. Traag, L. Waltman, N.J. van Eck, From Louvain to Leiden: guaranteeing well-connected communities, Scientific reports 9(1) (2019) 5233.

[43] A.D. Diehl, T.F. Meehan, Y.M. Bradford, M.H. Brush, W.M. Dahdul, D.S. Dougall, Y. He, D. Osumi-Sutherland, A. Ruttenberg, S. Sarntivijai, C.E. Van Slyke, N.A. Vasilevsky, M.A. Haendel, J.A. Blake, C.J. Mungall, The Cell Ontology 2016: enhanced content, modularization, and ontology interoperability, Journal of biomedical semantics 7(1) (2016) 44.

[44] V.Y. Kiselev, A. Yiu, M. Hemberg, scmap: projection of single-cell RNA-seq data across data sets, Nat Methods 15(5) (2018) 359–362.

[45] M.D. Robinson, D.J. McCarthy, G.K. Smyth, edgeR: a Bioconductor package for differential expression analysis of digital gene expression data, Bioinformatics (Oxford, England) 26(1) (2010) 139–40.

[46] L. Haghverdi, A.T.L. Lun, M.D. Morgan, J.C. Marioni, Batch effects in single-cell RNA-sequencing data are corrected by matching mutual nearest neighbors, Nature biotechnology 36(5) (2018) 421–427.

